# Interactive downstream proteomics analysis with MiraProt using Müller cell proteomes from equine recurrent uveitis

**DOI:** 10.64898/2026.08.27.747296

**Authors:** Adrian Schmalen, Amelie B. Fleischer, Barbara M. Riedel, Cornelia A. Deeg

## Abstract

Mass spectrometry-based proteomics requires downstream analysis of processed protein abundance data, including data inspection, filtering, statistical testing, functional enrichment, protein set comparison, network analysis, and visualization. MiraProt was developed as a modular, metadata-aware R Shiny platform that integrates these steps in a single interactive workflow for processed protein-level proteomics data. Its metadata-aware design enables identifiers, sample information, experimental conditions, transformations, and derived data columns to be defined during data preparation and reused consistently across downstream analyses.

To demonstrate its use, we reanalyzed a previously published label-free proteomic dataset of primary retinal Müller cells from healthy horses and horses with equine recurrent uveitis (ERU). ERU is a naturally occurring autoimmune eye disease of horses characterized by recurrent intraocular inflammation triggered by autoreactive T-cells. Müller cells are specialized retinal macroglia with various functions such as maintaining retinal ion homeostasis and supporting retinal neuron metabolism. Of 193 proteins with an adjusted p-value ≤ 0.05, 187 also showed at least a twofold difference in abundance between ERU-derived and control Müller cells. Functional enrichment highlighted nuclear RNA processing, chromatin-associated structures, DNA and RNA binding, interferon responses, and cell-cycle-associated programs. Gene set enrichment analysis identified positive enrichment of Interferon Alpha Response, Interferon Gamma Response, and MYC-, E2F-, and G2M-associated gene sets. Network analysis of shared proteins further linked this signature to DNA replication, mitotic checkpoint control, and RNA processing. ERU-derived Müller cells also showed increased abundance of the MHC class II protein HLA-DRA.

Together, these findings identified an interferon-responsive, cell-cycle-associated, and MHC class II-associated Müller cell protein signature in ERU and generated experimentally testable hypotheses for further mechanistic studies. MiraProt provides an accessible, metadata-aware framework for reproducible downstream exploration of processed proteomic datasets and prioritization of candidate proteins and pathways for experimental follow-up.

## Introduction

Mass spectrometry-based proteomics enables the quantitative analysis of thousands of proteins across biological conditions, but the interpretation of processed proteomic datasets remains challenging. After peptide identification, protein inference, and quantification by specialized mass spectrometry (MS) software, researchers often need to integrate quality control, statistical testing, functional enrichment, protein set comparison, network analysis, and visualization to extract biological meaning from the data (as reviewed in (1)). These downstream steps frequently require bioinformatics expertise and the use of multiple independent tools, which can complicate exploratory analysis, reduce reproducibility, and limit accessibility for users without programming experience (2–4).

MiraProt was developed to address this gap (5). It is a modular R Shiny-based platform for downstream analysis of processed protein-level proteomics data. The application complements existing mass spectrometry workflows rather than replacing raw-data processing software. MiraProt integrates data import, metadata assignment, filtering, imputation, batch correction, custom statistical comparisons, identifier conversion, quality control visualization, dimensionality reduction, volcano and dot plot visualization, Gene Ontology (GO) over-representation analysis, Gene Set Enrichment Analysis (GSEA), STRING-based protein association networks, Venn and UpSet analyses, and heatmap-based visualization within a single interactive environment. By combining established statistical and bioinformatic methods in a user-accessible workflow, MiraProt supports systematic exploration of proteomic datasets and facilitates the generation of potential biologically verifiable hypotheses.

To demonstrate the utility of MiraProt for hypothesis generation, we applied the platform to a previously published proteomic dataset of primary retinal Müller cells derived from healthy horses and horses affected by equine recurrent uveitis (ERU) (6). ERU is a naturally occurring inflammatory autoimmune eye disease of horses that shares important clinical and immunopathological features with human autoimmune uveitis (reviewed in (7, 8)). The disease exhibits a relapsing-remitting pattern with periods of acute inflammation followed by apparent clinical remission, closely mirroring autoimmune uveitis in humans. ERU pathogenesis involves CD4^+^ T cell-driven autoimmune responses targeting retinal autoantigens, among them interphotoreceptor retinoid-binding protein (9–12). The breakdown of immune privilege within the eye leads to infiltration of autoreactive lymphocytes (11, 13, 14), activation of granulocytes releasing Neutrophil Extracellular Traps (15) and subsequent tissue destruction, making ERU an invaluable translational model for understanding human uveal inflammatory diseases (9, 16, 17).

Retinal Müller cells, the predominant macroglial population, spanning the entire width of the retina, play central roles in both retinal homeostasis and pathology (18). In the healthy eye, they play an integral part in the inner blood retinal barrier (18), regulate pH, ion, and water balance (19–22), and facilitate neurotransmitter recycling including glutamate uptake and metabolism (23–25). In healthy retinas Müller cells do not proliferate with p27Kip1-mediated cell-cycle suppression (26, 27). During inflammatory conditions such as ERU, Müller cells undergo reactive gliosis characterized by morphological changes, upregulation of intermediate filaments (28, 29), proliferation (26, 30), and secretion of inflammatory mediators including cytokines and growth factors (31–33). This phenotypic transition represents a critical component of retinal inflammatory responses, though the underlying molecular mechanisms remain elusive.

The objective of this study was to present MiraProt and demonstrate its use for accessible, reproducible, and hypothesis-generating downstream analysis of processed proteomics data. Using ERU-derived Müller cells as a biologically relevant case study, we show how MiraProt supports quality assessment, differential abundance analysis, functional enrichment, protein set comparison, STRING-based network visualization, and heatmap-based exploration. This reanalysis illustrates how an integrated downstream analysis platform can help identify coherent biological signatures and prioritize candidate proteins and pathways for future mechanistic investigation.

## Materials and Methods

### Published proteomic dataset

This study is a secondary reanalysis of the previously published proteomic dataset generated by Fleischer et al. (6). The dataset comprises primary retinal Müller cell cultures derived from three healthy horses and three horses affected by equine recurrent uveitis (ERU), representing three biological samples per group. Healthy eyes were obtained from a local abattoir, whereas ERU eyes were obtained following therapeutic enucleation. ERU was diagnosed by veterinary ophthalmologists based on a history of at least three recurrent inflammatory episodes and characteristic clinical findings. Primary Müller cell isolation, culture, protein preparation, LC-MS/MS acquisition, and initial protein identification and quantification were performed as described previously (6).

Briefly, label-free LC-MS/MS measurements were acquired using a Q-Exactive HF-X mass spectrometer coupled to an Ultimate 3000 nano-RSLC system. Protein identification and quantification were performed with Proteome Discoverer 2.5 using Sequest HT and the Ensembl horse database (Release 75). The processed protein-level abundance matrix and protein annotations were used as the primary input for the present MiraProt reanalysis. Protein-level PSM counts were additionally used for DEqMS-based differential abundance analysis (34). The underlying mass spectrometry data are publicly available through PRIDE under accession PXD058170 (6).

### Ethics statement

The present study was based exclusively on reanalysis of previously published proteomic data and did not involve new animal experiments or new sample collection. The original equine eye samples were obtained from local abattoirs and cooperating equine clinics in accordance with the ethical principles and guidelines of the ARVO Statement for the Use of Animals in Ophthalmic and Vision Research. Collection of the samples was permitted by the local veterinary inspection office, Munich, Germany (permit number: DE 09 184 0063 21).

### MiraProt Software and Availability

MiraProt is a modular R Shiny application designed to facilitate comprehensive proteomic data analysis without requiring coding expertise. The application is based on the shiny web application framework (version 1.14.0) and the R packages shinyjs (version 2.1.1) and shinyWidgets (version 0.9.1) (35–37). For visualization, plots rely on the R package ggplot2 (version 4.0.3) and plotly (version 4.12.1) (38, 39). Interconnected modules operate on a shared processed dataset and its associated metadata, allowing analysis outputs and protein selections to be reused across downstream modules. The MiraProt source code is openly available under the MIT License at GitHub (https://github.com/AdSchmalen/MiraProt, accessed on 8/27/2026). Analyses were performed using MiraProt version 1.0.0 and preceding development builds. The finalized MiraProt version 1.0.0 used as the reference implementation for this study is archived on Zenodo (DOI: 10.5281/zenodo.22110860) (5). Package dependencies and their versions for the MiraProt v1.0.0 reference environment are recorded in an renv.lock file included with the archived software release, enabling restoration of the R package environment using renv (package version 1.2.4).

### Data Wizard

The Data Wizard serves as the central preprocessing and data-definition interface of MiraProt. It imports, annotates, restructures, processes, and exports protein-level proteomics datasets for downstream analysis. Because outputs from mass spectrometry software vary in structure and terminology, the Data Wizard uses a flexible metadata system to define the analytical role of individual columns, including protein identifiers, abundance values, abundance ratios, statistical parameters, confidence information, and annotation columns.

Datasets can be imported in XLSX, CSV, or TSV format, with spreadsheet-specific selection of sheets and header rows. The primary requirement for downstream analysis is one protein per row; long- and wide-format tables can be interconverted, and additional tables can be merged using user-defined key columns. The resulting processed table and its associated metadata are shared across downstream MiraProt modules.

Metadata link abundance columns to experimental samples and conditions and associate ratio columns with their corresponding numerator and denominator conditions. Multiple identifier columns can be retained for compatibility with different analyses and external databases. Metadata assignment can be performed manually or assisted by regular-expression-based header matching, and assignment rules can be stored as reusable templates. A predefined template for Proteome Discoverer exports was used as the starting point for the dataset analyzed here.

Preprocessing options include protein-confidence and valid-value filtering, custom numeric or categorical filters, data transformations, and replacement of user-defined values. MiraProt additionally provides optional batch- and intensity-correction methods, including ComBat, limma-based correction, LOESS, offset, and quantile correction, as well as left-censored, Random Forest, and MICE/CART-based missing-value imputation (40–42). Minimum valid-value requirements can be defined for statistical comparisons.

Custom pairwise contrasts can be calculated directly from abundance values using user-defined numerator and denominator groups. MiraProt calculates corresponding summary statistics, abundance ratios, p-values, adjusted p-values, and optional basemean values. Available statistical methods include Student’s and Welch’s t-tests, moderated Welch testing, limma, DEqMS, and the Mann-Whitney U test (34, 43–46), with several multiple-testing correction procedures available (47–52). DEqMS additionally uses protein-level peptide-spectrum-match or peptide-count information for variance estimation (34).

Identifier harmonization is supported through AnnotationHub/OrgDb for within-species mapping and BioMart/Ensembl for intra- and inter-species mapping (53, 54). Supported identifiers include commonly used gene, Ensembl, Entrez, and UniProt identifiers, with additional identifier types available depending on the selected annotation backend. Identifiers can be retained individually or merged into a combined identifier column.

Processed datasets and analysis outputs can be exported as multi-sheet Excel workbooks. The comprehensive export contains the original and processed data, associated metadata, derived analysis columns, and outputs from available downstream modules, including GO/GSEA, heatmap, Venn/UpSet, PCA, Volcano, DotPlot, and STRING analyses. Log and summary sheets additionally document the exported dataset and analysis session.

### Dataset preprocessing and differential abundance analysis

Human orthologue gene symbols were used for downstream functional analyses. Missing, ambiguous, locus-based, numeric, uncharacterized, or otherwise non-standard gene annotations were manually curated where possible or mapped to human gene symbols using BioMart based on equine accession identifiers. Original and BioMart-derived annotations were merged while prioritizing existing gene symbols, and the resulting Merged_ID_first_2cols identifier was used for downstream analyses.

Proteins with fewer than three quantified abundance values across the six samples or fewer than two unique peptides were excluded. A duplicate HNRNPD mapping was resolved by retaining ENSECAP00000007637 and removing the uncharacterized entry ENSECAP00000020212. Proteins annotated as keratins were also excluded. After filtering, 3,776 proteins remained for downstream analysis.

Missing normalized abundance values were imputed using left-censored imputation. Differential protein abundance between ERU-derived Müller cell samples and healthy controls was assessed using a custom DEqMS contrast (34), with protein-level peptide-spectrum-match counts used for variance estimation. P-values were adjusted for multiple testing using the Benjamini-Hochberg false discovery rate procedure (51). A basemean column representing the mean normalized abundance across all six samples was additionally calculated.

Proteins were considered differentially abundant at an adjusted p-value ≤ 0.05 and an absolute twofold abundance difference. Accordingly, ERU/healthy log_2_ fold change ≥ 1 were classified as increased in ERU-derived Müller cells and log_2_ fold change ≤ -1 as decreased.

Data processing and automated identifier mapping were performed using MiraProt’s Data Wizard. Manual annotation curation was performed as described above.

### Data quality assessment modules

MiraProt provides several modules for quality assessment and exploratory analysis of processed proteomics data. These modules support inspection of the processed data matrix, assessment of sample-wise abundance distributions and protein identification patterns, and exploration of sample or protein relationships by dimensionality reduction.

The table module (“Table”) displays the current processed data matrix used by downstream MiraProt modules. The abundance module (“Abundances”) visualizes sample-wise abundance distributions using box plots and can additionally be used to assess distributional changes following missing-value imputation. The sample identification module (“Sample IDs”) summarizes protein identifications per sample using metadata-annotated sample- or file-detection columns and supports visualization of categorical identification patterns as stacked bar plots. For the dataset analyzed in this study, entries classified as “High” represent proteins directly identified from MS2 spectra, whereas “Peak Found” represents protein entries quantified by feature matching without direct MS2 identification in the respective sample.

For exploratory analysis of high-dimensional proteomics data, MiraProt implements principal component analysis (PCA) (55, 56) and uniform manifold approximation and projection (UMAP) (57) for either samples or proteins. Before dimensionality reduction, duplicate protein identifiers are made unique and constant variables are excluded. Selected abundance values are retransformed according to their metadata where applicable, log_2_-transformed when required, and standardized protein-wise across samples. For PCA of the non-imputed dataset, proteins with missing normalized abundance values were excluded. PCA after imputation was performed using the left-censored-imputed abundance values. The PCA is performed using stats::prcomp(), with explained variance calculated from the squared component standard deviations. MiraProt additionally supports UMAP through the umap R package (version 0.2.10.0), with configurable neighborhood, distance, and minimum-distance parameters (57).

### Data visualization modules

MiraProt includes visualization modules for graphical exploration of quantitative protein-level data, including differential abundance and comparison-specific protein patterns.

Volcano plots are generated from abundance-ratio columns and corresponding p-value or adjusted p-value columns. The x-axis represents the log_2_ abundance ratio and the y-axis the negative log_10_-transformed significance value. Proteins are classified as increased, decreased, or not differentially abundant according to user-defined significance and fold-change thresholds.

For datasets containing multiple comparisons, MiraProt pairs abundance-ratio and significance columns primarily using metadata annotations defined in the Data Wizard, with pattern-based column-name matching available as a fallback. Separate volcano plots are generated for matched comparisons. Static plots are generated using ggplot2 with optional labeling through ggrepel, whereas interactive plots use plotly and provide protein identifiers and plot coordinates.

A generalized DotPlot module additionally supports scatter-, volcano-, and MA-like visualization of numeric protein-level variables (58).

### Functional enrichment analysis

Functional interpretation in MiraProt is supported by modules for Gene Ontology (GO) over-representation analysis (ORA) and Gene Set Enrichment Analysis (GSEA). These modules analyze user-defined protein subsets or ranked protein lists and allow resulting protein sets to be transferred to other MiraProt modules for downstream visualization and comparison.

GO ORA is performed on subsets selected from the processed analysis table. Species-specific annotations are retrieved through AnnotationHub or locally available OrgDb resources, with user-defined identifier columns and key types. Missing or empty identifiers are removed and duplicate identifiers collapsed before enrichment. Unmapped identifiers are excluded. A custom background universe can be supplied; otherwise, all valid identifiers in the selected identifier column of the processed dataset are used as the reference universe.

GO enrichment is performed using clusterProfiler::enrichGO() (clusterProfiler version 4.20.0) (59–62). Molecular Function, Cellular Component, and Biological Process can be analyzed individually or jointly. P-value and q-value thresholds, multiple-testing correction, and gene-set size limits are user-configurable. Benjamini-Hochberg correction is the default. Pairwise term similarity is calculated using pairwise_termsim() to support enrichment-map visualization.

The GSEA module analyzes ranked protein lists generated either from user-defined group comparisons or from precomputed quantitative and statistical variables (63–65). Multiple ranking approaches are available, including fold-change-, significance-, and group-based statistics. Rankings can be generated from original or imputed abundance values where available, and optional tie-breaking can be applied to ambiguous ranks.

GSEA uses local GMT gene-set files, including MSigDB-derived or user-defined collections, and is performed using clusterProfiler::GSEA() with the fgsea backend (59–62, 66). The number of permutations and gene-set size limits are configurable, and p-values are adjusted using Benjamini-Hochberg correction. Optional PADOG-style weighting can be applied to reduce the influence of proteins represented across many gene sets (67). GO and GSEA results can be visualized using complementary enrichment plots, and proteins associated with selected GO terms or GSEA core-enrichment sets can be transferred to downstream MiraProt modules.

For the present study, GO ORA was performed using human gene symbols from the Merged_ID_first_2cols column. Proteins were selected using a log_2_ fold-change cutoff > 1 and an adjusted p-value < 0.05 based on the DEqMS ERU-versus-control contrast. Molecular Function, Biological Process, and Cellular Component were analyzed using *Homo sapiens* annotations with SYMBOL as key type together with the Merged_ID_first_2cols gene symbols. P-values were adjusted using the Benjamini-Hochberg method; the p-value cutoff was 0.05, q-value cutoff 0.2, and minimum and maximum gene-set sizes were 10 and 500, respectively. No custom background universe was supplied. Thus, all valid human gene symbols in the processed dataset were used as the enrichment background.

GSEA was performed using the Merged_ID_first_2cols gene symbols and the Hallmark collection h.all.v2025.1.Hs.symbols.gmt (68). Rankings were generated from normalized, left-censored imputed abundances comparing ERU-derived Müller cells with healthy controls using the Fold Change Rank Ordering Statistics method (65). GSEA was performed with 1,000 permutations, a gene-set size range of 10–500, and Benjamini-Hochberg multiple-testing correction. Ties were resolved randomly. The MiraProt minimum-valid-values setting was 1 with the validation rule set to “In total.”

### Protein set and network analysis

STRING-based protein association networks are generated using the STRING database through the STRINGdb R package (version 2.24.0) (69). Unmapped proteins are excluded and duplicate identifiers collapsed before network construction. Networks can optionally be expanded by adding STRING-supported neighboring proteins not present in the original input set.

Networks are rendered interactively using visNetwork (version 2.1.4) (70). Topological communities are identified using edge-betweenness clustering implemented in igraph (version 2.3.3) (71–73) and are used to support interpretation of network structure.

For the present analysis, protein interaction networks were generated from the nine core-enriched proteins shared by the HALLMARK_MYC_TARGETS_V1, HALLMARK_E2F_TARGETS, and HALLMARK_G2M_CHECKPOINT gene sets. Networks were constructed using STRING database version 12.0 with *Homo sapiens* as organism, a confidence score threshold of 700, and the interaction setting “Full”. Networks were analyzed either without additional interactors or after addition of the two first-order neighboring proteins with the highest number of connections to the seed network.

Protein-set overlaps can be visualized using Venn diagrams or UpSet plots (74), implemented with the VennDiagram package (version 1.8.2) (75) and ComplexUpset (version 1.3.3) (74, 76), respectively.

Heatmaps were generated using ComplexHeatmap (version 2.28.0) (77). For the present analysis, the 29 proteins occurring in at least two of the MYC Targets V1, E2F Targets, and G2M Checkpoint gene sets were visualized using normalized, left-censored-imputed abundance values. Abundance values were log_2_-transformed and standardized protein-wise to row-wise z-scores for visualization. Sample order was determined by hierarchical clustering of the log_2_-transformed abundance matrix using Euclidean distance and Ward’s D2 linkage. Pairwise protein correlations were calculated using Pearson correlation on the row-wise standardized expression matrix, and protein ordering was based on the resulting correlation structure. Additional annotation tracks displayed log_2_ basemean and log_2_ ERU/control abundance ratios.

### Use of generative AI-assisted tools

Generative AI-assisted tools were used during development of MiraProt and preparation of the manuscript. OpenAI ChatGPT models used during the project included GPT-5.4, GPT-5.5, and 5.6. Anthropic Claude models used during the project included Claude Sonnet 4.5/4.6, and Claude Opus 4.5/4.6.

These tools were used for AI-assisted code development, troubleshooting, refactoring, and debugging of R/Shiny code, as well as for language editing, restructuring of manuscript sections, and improvement of clarity and readability. All AI-assisted code suggestions were reviewed, modified where necessary, and tested by the authors before inclusion in MiraProt. All manuscript text, analyses, interpretations, and conclusions were critically reviewed and approved by the authors. The authors take full responsibility for the final content of the manuscript and software. No generative AI tool was listed as an author.

## Results

We reanalyzed a previously published proteomic dataset using MiraProt. The dataset contained protein abundance measurements from six Müller cell samples (three ERU-affected, three healthy controls). The original data contained 5,232 identified proteins. Initial quality filtering retained 3,776 proteins for downstream statistical analysis. Pre-imputation abundance distributions were similar across samples, with median log_2_-transformed abundance values ranging from 20.20 (sample F7_C) to 20.47 (sample F3_ERU). The interquartile ranges (Q1-Q3) were comparable (Fig 1A), indicating comparable abundance distributions and no obvious global sample shift.

**Fig 1.**
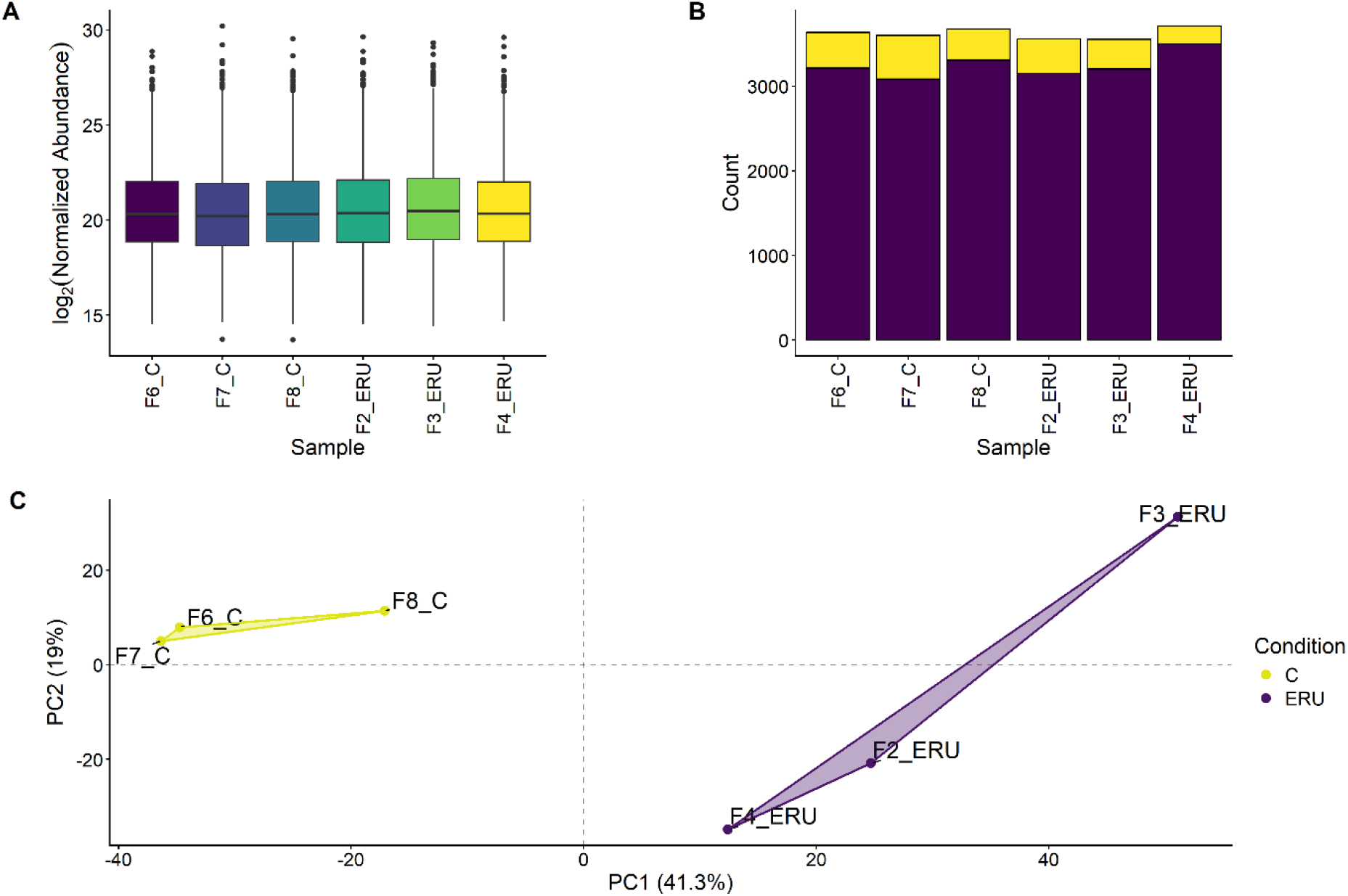
Quality control of MS results. **(A)** Boxplot summary of normalized abundance values identified in the different samples. **(B)** Staggered bar plot with protein identifications per sample. Purple bars (“High”) represent proteins directly identified from MS2 spectra, whereas yellow bars (“Peak Found”) represent protein entries quantified by feature matching without direct MS2 identification in the respective sample. **(C)** Principal component analysis (PCA) of the samples showing the two main principal components PC1 (x-axis) and PC2 (y-axis). ERU data points are depicted in purple and connected by a purple polygon and control data points C are depicted in yellow connected by a yellow polygon.

Left-censored imputation replaced 1,295 missing values across the six samples and shifted sample medians slightly toward lower abundance values. Post-imputation, median log_2_ abundance values decreased to 19.97-20.23, reflecting the addition of low-abundance imputed values (S1 Fig A).

Protein identification efficiency varied modestly across samples. Sample F4_ERU showed the highest number of identified proteins (3,712 total: 3,499 by direct MS2 detection, 213 by match-between-runs), while sample F3_ERU had the lowest (3,553 total: 3,202 by MS2 detection, 351 by match-between-runs). Direct MS2-based identifications were lowest in sample F7_C (3,601 total: 3,082 proteins by MS2 detection, 519 by match-between-runs) (Fig 1B). Protein identification numbers were comparable across samples, with modest variation in the relative contribution of direct MS2 detection and match-between-runs identifications.

Principal component analysis showed clear separation between ERU-affected Müller cell samples and healthy controls. The first principal component (PC1) explained 41.3% of the total variance, while PC2 accounted for 19.0% of the variance. Samples derived from ERU Müller cells exhibited exclusively positive PC1 values, whereas controls showed exclusively negative PC1 values, consistent with condition-associated differences in the proteomic profiles. Control samples formed a tight cluster with minimal variance along PC2, while ERU samples displayed greater heterogeneity, with PC2 values ranging from -34.8 to 31.3 (Fig 1C). This pattern suggests that ERU-affected samples exhibit a broader inter-individual variability in protein abundance profiles compared to healthy controls in this dataset. The condition-associated separation was retained after left-censored imputation (S1 Fig B).

Of the 193 proteins with adjusted p-value ≤ 0.05, 187 also met a twofold abundance-change threshold (|log2FC| ≥ 1), including 145 proteins with increased abundance in ERU samples and 42 proteins with decreased abundance. Log_2_ abundance ratios ranged from -7.20 to 6.99 (Fig 2A; S1 Table). Among the proteins with increased abundance in ERU-derived Müller cells, the MHC class II-associated protein HLA-DRA was exclusively detected in Müller cells derived from ERU affected horses with direct MS2 detections and four unique peptides, as already pointed out in the original publication of this dataset (6).

**Fig 2.**
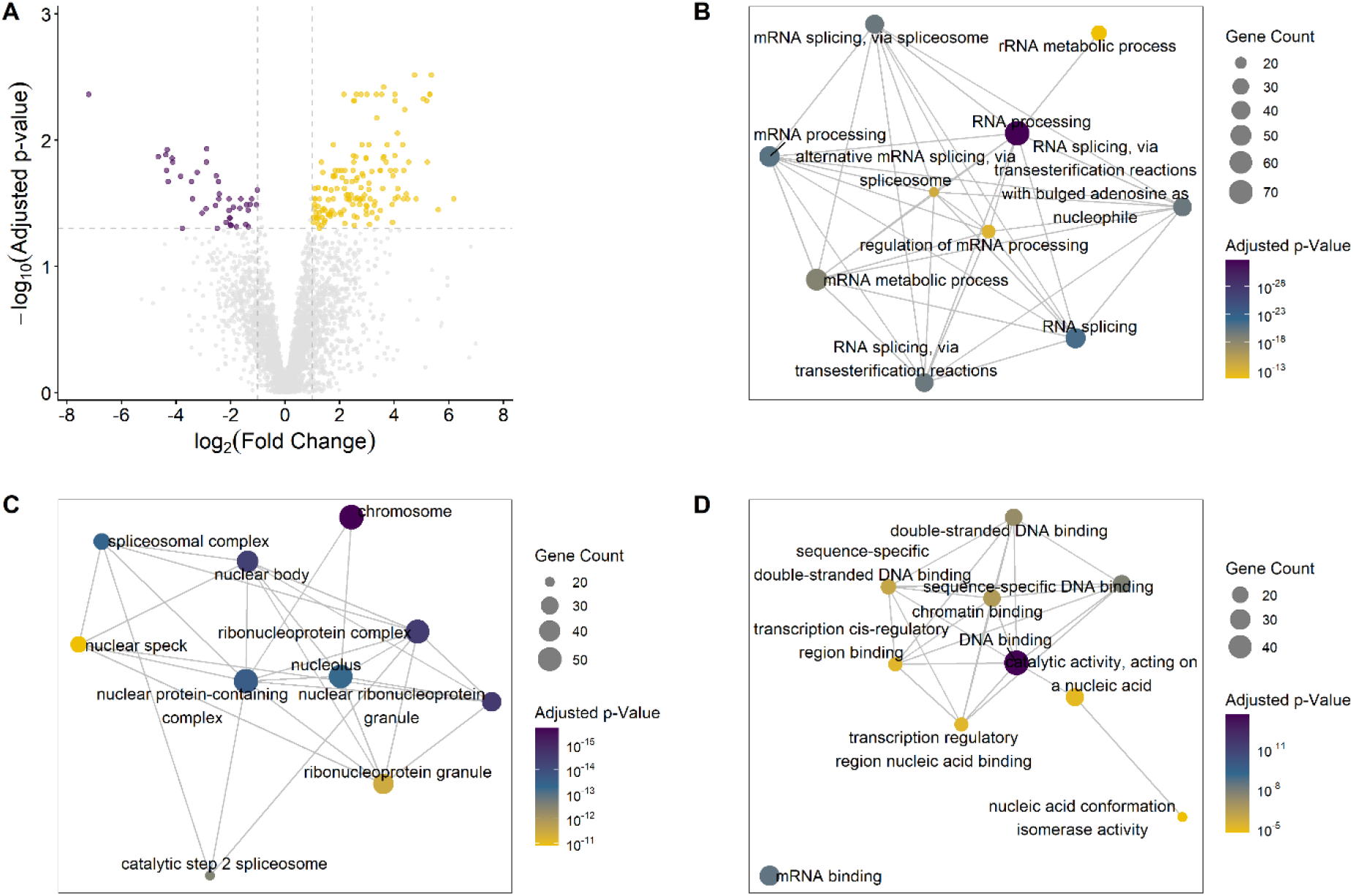
GO Enrichment Analysis. Volcano plot showing differentially abundant proteins in retinal Müller cells derived from ERU cases compared with control Müller cells. Proteins with significantly increased abundance are shown in yellow (log_2_ fold change ≥ 1, adjusted p value ≤ 0.05). Proteins with significantly decreased abundance in ERU-derived Müller cells relative to controls are shown in purple (log_2_ fold change ≤ −1, adjusted p value ≤ 0.05) **(A)**. Gene Ontology enrichment analysis was performed for proteins significantly more abundant in ERU Müller cells compared to control. Enrichment maps display the ten most significant GO terms for the domains biological process **(B)**, cellular component **(C)**, and molecular function **(D)**. Dot size reflects the number of associated genes, and color indicates the adjusted p value. GO terms sharing common proteins are connected by edges.

GO enrichment analysis was performed on the 145 proteins meeting both criteria (adjusted p-value ≤ 0.05, log₂ abundance ratio ≥ 1). The analysis covered three GO domains: Biological Process (BP), Cellular Component (CC), and Molecular Function (MF). 202 GO terms across the three GO domains were significantly enriched with an adjusted p-value < 0.05. Enrichment maps of the top 10 terms per domain showed extensive overlap between related GO terms, reflecting shared contributing proteins among enriched categories.

In the Biological Process domain, enriched terms were predominantly associated with RNA processing and splicing mechanisms (Fig 2B). The Cellular Component analysis revealed significant enrichment in nuclear structures, with the top enriched terms including chromatin, nuclear speck, fibrillar center, and nucleolus (Fig 2C). Molecular Function analysis identified DNA and RNA binding activities as the most significantly enriched categories (Fig 2D). The GO analysis highlighted enrichment of nuclear RNA-processing-related terms in Müller cell cultures from ERU-affected eyes compared to healthy controls.

GSEA was performed using the Hallmark gene set collection from MSigDB, comparing ERU samples to control samples (68). Protein rankings were derived using the Fold Change Rank Ordering Statistics algorithm applied to left-censored, imputed, and normalized abundance values. GSEA was performed with 1,000 permutations using the settings described above.

GSEA identified 15 Hallmark gene sets with significant enrichment at an adjusted p-value ≤ 0.05 (Fig 3). Seven of these gene sets were positively enriched, as indicated by a positive normalized enrichment score (NES). These gene sets are primarily associated with cell-cycle regulation and proliferative signaling, including E2F Targets (NES = 2.30, adjusted p-value = 8.82 × 10^-10^, mapped gene set size = 77), G2M Checkpoint (NES = 2.29, adjusted p-value = 5.76 × 10^-8^, mapped gene set size = 62), Myc Targets V1 (NES = 2.21, adjusted p-value = 7.20× 10^-11^, mapped gene set size = 165).

**Fig 3.**
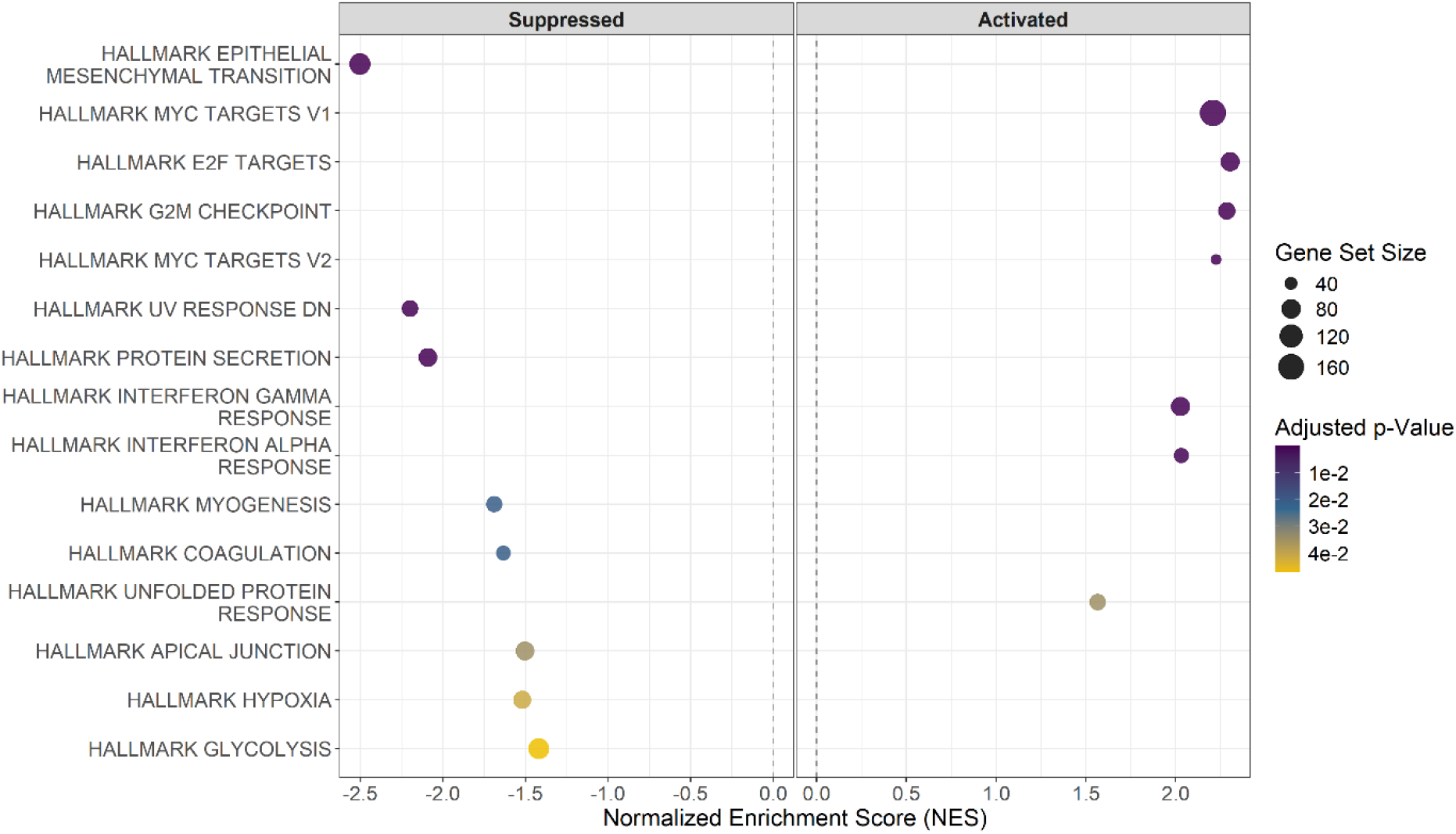
GSEA of ERU-affected Müller cells compared to healthy controls. Enrichment score dotplot of significantly enriched GSEA gene sets of h.all.v2025.1.Hs.symbols.gmt. Dot size represents mapped gene set size. Color indicates the adjusted p-values of the identified enriched gene sets.

In addition, immune- and stress-related gene sets show positive enrichment, including Interferon Gamma Response (NES = 2.03, adjusted p-value = 1.10 × 10^-5^, mapped gene set size = 74), Interferon Alpha Response (NES = 2.03, adjusted p-value = 7.83 × 10^-5^, mapped gene set size = 48), and the Unfolded Protein Response.

Eight gene sets are negatively enriched in Müller cells derived from ERU samples. The most pronounced negative enrichment occurs in Epithelial Mesenchymal Transition (NES = -2.50, adjusted p-value = 7.20 × 10^-11^, mapped gene set size = 95), UV Response DN (NES = -2.20, adjusted p-value = 7.89 × 10^-6^, mapped gene set size = 56), and Protein Secretion (NES = -2.09, adjusted p-value = 1.10 × 10^-5^, mapped gene set size = 72). Additional negatively enriched gene sets include Glycolysis, Hypoxia, Myogenesis, Xenobiotic Metabolism and Coagulation (S1 Table, Sheet 5_GSEA_Analysis).

Myc Targets V1 contributed the largest number of core-enriched proteins among the positively enriched Hallmark sets, comprising 96 core-enriched proteins, of which 72 are unique to this set (Fig 4A). Core-enriched proteins correspond to the leading-edge subset. The Interferon Gamma Response (37 proteins) and Interferon Alpha Response (26 proteins) exhibit the greatest overlap, sharing 20 proteins.

**Fig 4.**
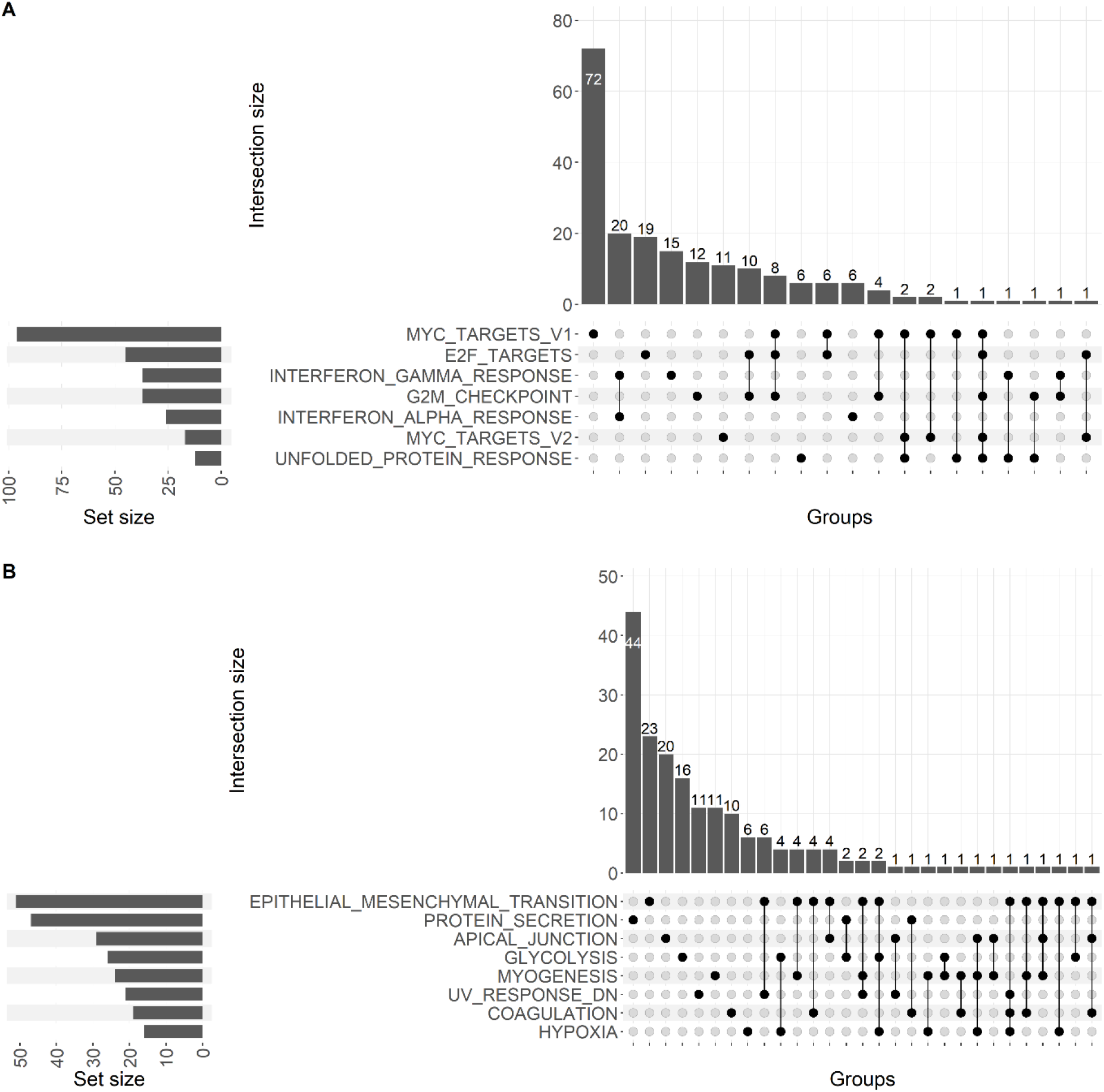
Protein overlaps among significantly enriched GSEA gene sets. UpSet plots summarizing overlaps among positively **(A)**, and negatively **(B)** enriched gene sets. Each column in the dot matrix represents a specific intersection. Connected dots indicate the gene sets included in that intersection. Vertical bars denote the number of proteins shared among the indicated sets (intersection size). Horizontal bars indicate the total number of core enriched proteins associated with each gene set. Only significantly enriched gene sets (Benjamini-Hochberg adjusted p-value < 0.05) are shown. Intersections are based on core-enriched proteins.

A smaller intersection was observed among Myc Targets V1, E2F Targets, and G2M Checkpoint, with nine proteins common to all three sets. Collectively, these three gene sets contain 140 core enriched proteins, including 111 unique proteins and 29 present in at least two gene sets. Together, these gene sets represent cell-cycle-associated programs linked to growth, DNA replication, and mitotic progression (68, 78, 79). Despite related gene set names, Myc Targets V1 and Myc Targets V2 shared only five core-enriched proteins in this dataset.

NOLC1 is the most recurrent protein among positively enriched Hallmark gene sets, appearing in five of the seven sets. Notably, it is absent only from the interferon response gene sets.

Negatively enriched gene sets showed less overlap than positively enriched gene sets. The largest overlap comprised six proteins shared between Epithelial Mesenchymal Transition and UV Response DN, whereas most other intersections contained one to four proteins (Fig 4B). Among negatively enriched sets, Epithelial Mesenchymal Transition contributed the largest number of core-enriched proteins in control Müller cells (95 proteins), followed by Protein Secretion (72 proteins).

To further assess abundance patterns within the proliferation-associated GSEA overlap, we analyzed the 29 proteins present in at least two of the three enriched gene sets Myc Targets V1, E2F Targets, and G2M Checkpoint. Sample ordering based on Euclidean distance visually separated control Müller cell proteomes from ERU-derived Müller cell proteomes. Across this protein set, ERU-derived Müller cell samples showed predominantly positive row-wise z-scores, whereas control samples showed predominantly negative row-wise z-scores, indicating higher relative abundance of these proteins in ERU-derived samples. This was expected since proteins were chosen from positively enriched gene sets.

Protein ordering was based on Pearson correlation of abundance profiles across samples. The nine proteins shared by all three gene sets were highlighted by gene symbol to distinguish them from proteins present in two of the three gene sets. The nine proteins shared by all three cell-cycle-associated gene sets were TRA2B, SYNCRIP, SRSF1, HNRNPD, MCM6, MCM2, MAD2L1, KPNA2, and NOLC1. Among the shared proteins, SRSF1, HNRNPD, and TRA2B showed similar abundance profiles and formed a correlation-defined subgroup in the protein correlation heatmap. In contrast, the remaining proteins shared by all three gene sets were distributed across the heatmap rather than forming a single correlation-defined block. Proteins in the analyzed overlap showed positive log_2_ abundance ratios and log_2_ basemean values ranging from 16 to 24, indicating that the proteins were detected across a broad abundance range and were increased in ERU-derived samples (Fig 5A).

**Fig 5.**
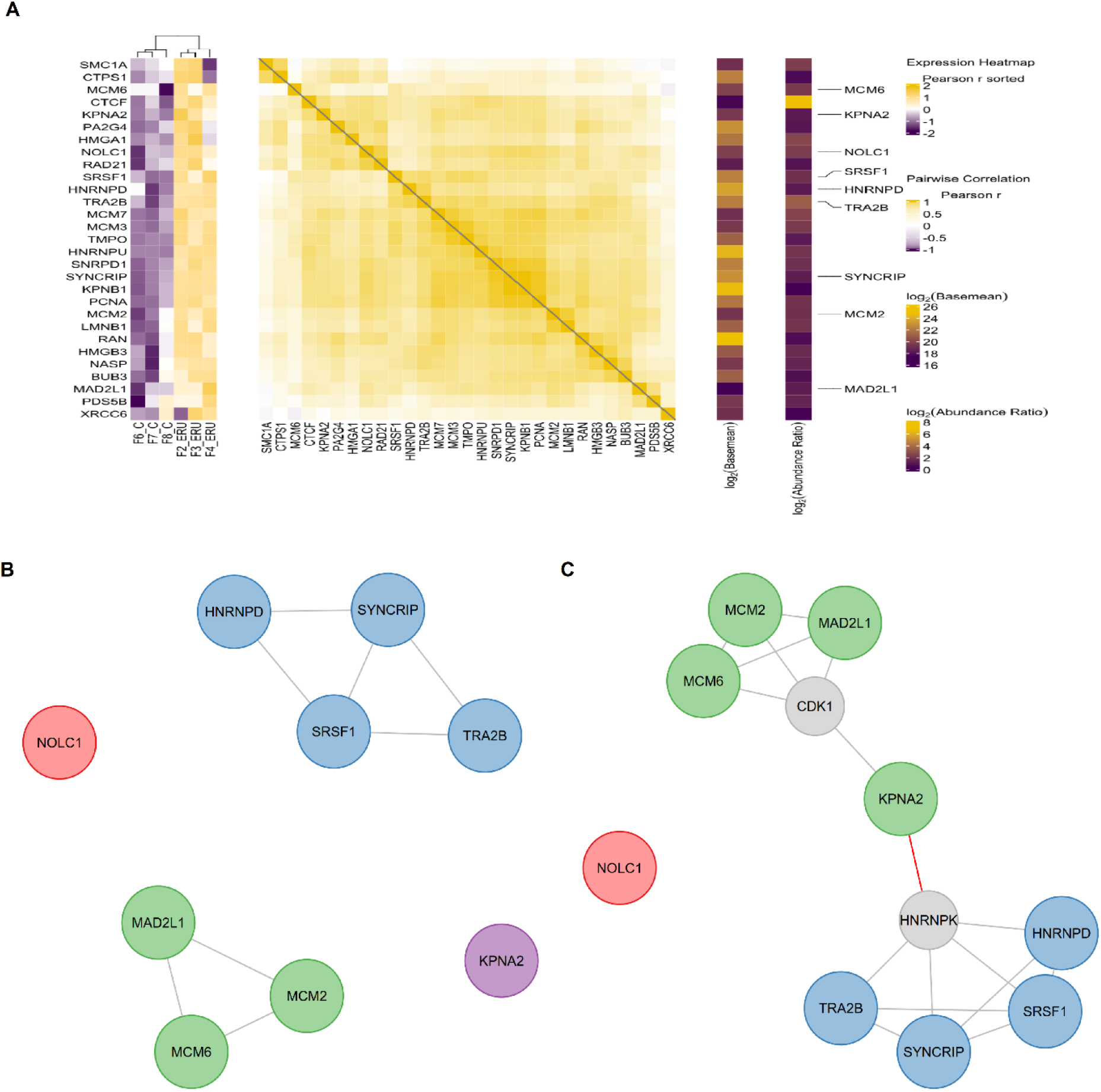
Core enriched proteins of proliferation-associated gene sets. **(A)** Heatmap-based analysis of proteins present in at least two of the three enriched gene sets Myc Targets V1, E2F Targets, and G2M Checkpoint. Proteins present in all three gene sets have been highlighted with their gene symbol on the right. The panel shows the z-score-based abundance heatmap, the protein correlation heatmap, and single-column annotation heatmaps for log_2_ basemean and log_2_ abundance ratio. In the z-score-based heatmap components, columns correspond to samples and rows correspond to proteins. Sample ordering was determined by hierarchical clustering of log_2_-transformed, normalized, and imputed abundance values using Euclidean distance and Ward’s D2 linkage. Protein-wise z-scores were calculated from the log_2_-transformed abundance matrix. Pairwise protein correlations were calculated using Pearson correlation on the row-wise standardized expression matrix, and protein ordering was based on the resulting correlation structure. **(B)** STRING interaction network of the core enriched proteins from Myc Targets V1, E2F Targets, and G2M Checkpoint. **(C)** STRING network after inclusion of two first-order neighboring proteins. Edges represent known and predicted functional and physical interactions. Red edges highlight inter-cluster connections. Node colors denote cluster membership. Grey nodes indicate added neighboring proteins that were not part of the core enriched protein set. Networks were generated using STRING database v12.0 with a confidence score threshold of 700.

Among the positively enriched gene sets, Myc Targets V1, E2F Targets, and G2M Checkpoint represented the main cell-cycle-associated sets and, with 29 proteins occurring in at least two of the three gene sets, showed substantial overlap. To further characterize the most stringent overlap at the network level, we performed STRING network analysis on the nine proteins shared by all three gene sets incorporating both physical and functional interactions. This analysis revealed two connected multi-protein clusters. One comprising TRA2B, SYNCRIP, SRSF1, and HNRNPD, and a second containing MCM6, MCM2, and MAD2L1 (Fig 5B). SYNCRIP and SRSF1 each formed three edges, representing the highest degree among the connected seed proteins.

KPNA2 and NOLC1 did not display direct interactions with the other shared proteins in this network. To examine whether the two modules could be connected through highly connected STRING-supported intermediary proteins, the two first-order neighbors with the highest number of edges to the seed network were added. This expansion connected the two modules through KPNA2, with CDK1 and HNRNPK acting as intermediary nodes (Fig 5C). HNRNPK (Accession ENSECAP00000022686) was detected in the proteomic dataset but was not differentially abundant between ERU and control samples (p = 0.34; adjusted p = 0.75). CDK1 was not detected in the proteomic dataset. Among the seed proteins, SRSF1 and SYNCRIP had the highest number of edges with four edges each, suggesting a central topological position among the proteins shared by the three cell-cycle-associated gene sets. Although NOLC1 was part of five of the seven positively enriched GSEA gene sets, it did not connect to the shared Myc Targets V1, E2F Targets, and G2M Checkpoint proteins in the STRING network, even after first-order neighbor expansion.

Müller cells derived from ERU-affected horses displayed signatures of enhanced nuclear RNA processing, interferon responsiveness, and cell-cycle-associated activity compared to healthy controls. Integration of enrichment and network analyses identified nine proteins shared across the Myc Targets V1, E2F Targets, and G2M Checkpoint gene sets, representing a core component of the proliferation-associated protein signature.

## Discussion

MiraProt was applied in this study for quality assessment and downstream reanalysis of a previously published proteomic dataset (6). It integrates filtering, visualization, differential abundance analysis, and enrichment analysis within a single framework (34, 41–46, 57, 61, 63, 65, 69, 74, 76, 80–83). A practical advantage of this integration is the metadata-aware handling of processed proteomic data: identifiers, sample information, experimental conditions, transformations, and derived data columns can be defined during data preparation either manually or automatically and subsequently reused across multiple downstream analyses. This reduces the need for repeated restructuring and manual transfer of intermediate data between separate analytical tools and facilitates consistent analysis of the same dataset across complementary approaches. In the present study, this framework enabled systematic reanalysis of the processed dataset and supported the identification of molecular patterns that were not the primary focus of the original analysis (6). The present reanalysis therefore provides a proof-of-principle application of MiraProt for its intended purpose without implying replacement of the established statistical and bioinformatic methods implemented by the underlying analytical packages. For longer-term use, MiraProt is openly available under the MIT License, with source code hosted on GitHub and versioned releases archived on Zenodo. Its modular architecture, reproducible dependency environment, and session save/restore functionality support maintainability, future extension, and reproducible reuse.

P-values reported by primary mass spectrometry analysis software, such as Proteome Discoverer, may differ from those obtained using downstream tools such as MiraProt, because differential abundance results depend on multiple workflow steps, including raw data quantification, abundance matrix construction, normalization, missing-value handling, and the statistical model used for inference (84, 85). This discrepancy is expected because proteomics workflows can operate at different levels of data processing, ranging from peptide-level models to analyses based on summarized protein-level abundance values (85, 86). Primary proteomics workflows may incorporate peptide-level and, depending on the workflow, spectrum- or PSM-level information during quantification, protein summarization, variance estimation, and statistical testing (34, 86). In contrast, MiraProt operates on processed protein-level abundance values and applies statistical tests directly to summarized data, similar in principle to other downstream protein-level analysis tools (87, 88). This approach is transparent and reproducible when processed datasets and analysis settings are available, but it does not explicitly model peptide-level variability or uncertainty introduced during protein inference (34, 85, 86). For this reason, the present analysis was used to explore additional biological patterns in the processed dataset rather than to benchmark the original raw-file-based statistical output.

The present reanalysis should be viewed as an extension of the original study rather than as a replacement of its primary analysis. The original publication focused on differential protein abundance in ERU-derived Müller cells and identified ARG1, MRC2, THBS1, and MHC class II-associated proteins as markers of disease-associated changes in immune-related Müller cell functions (6). In agreement with this interpretation, the MiraProt-based reanalysis retained an inflammatory and interferon-associated component, consistent with the immune phenotype described previously. The main extension of the present analysis was the systematic evaluation of the processed protein-level dataset by GSEA, GO enrichment, protein set overlap analysis, and STRING-based association networks. This shifted the analytical focus from individual marker proteins toward coordinated molecular programs and revealed an additional proliferation-associated and RNA-processing signature in ERU-derived Müller cells. Thus, the reanalysis does not contradict the original findings but places them into a broader pathway-level context and nominates additional candidates for experimental follow-up.

The reanalysis revealed a consistent proliferation-associated molecular signature in Müller cells derived from ERU-affected horses. Four enriched Hallmark gene sets supported this interpretation, including Myc Targets V1, Myc Targets V2, E2F Targets, and G2M Checkpoint, together with GO terms related to DNA replication and chromosome organization (63, 68, 89). Although c-Myc itself was not directly identified as differentially abundant in this dataset, enrichment of Myc target gene sets suggests activation of c-Myc-associated transcriptional programs. These gene sets cover related aspects of cell-cycle-associated regulation: MYC-associated programs are linked to cellular growth, metabolism, biosynthetic activity, and proliferation (90, 91). E2F target genes regulate cell-cycle entry and progression, particularly the G1/S transition (79, 92, 93). The G2M Checkpoint genes are associated with preparation for mitosis and maintenance of genomic integrity (68, 94). The overlap of Myc Targets V1, E2F Targets, and G2M Checkpoint comprised nine proteins, indicating that part of the enriched signal was shared across multiple proliferation-associated programs. While this is not direct proof of proliferation, it may reflect gliotic activation, proliferative priming, or stress-associated remodeling rather than active cell division alone. Validation by independent markers such as Ki-67, PCNA, EdU incorporation, or cell-cycle profiling would be required to determine whether this molecular signature corresponds to increased proliferative activity.

STRING DB analysis added functional detail to this proliferation-associated signature. One module, containing MCM2, MCM6, and MAD2L1, links the enrichment result to DNA replication licensing and mitotic checkpoint control (95, 96). MCM2 and MCM6 are components of the MCM2-7 helicase complex, whereas MAD2L1 is involved in spindle assembly checkpoint regulation (95, 96). This combination points to a cell-cycle-associated program that includes both preparation for genome duplication and control of chromosome segregation. A second module consisted of TRA2B, SRSF1, SYNCRIP, and HNRNPD, all of which are linked to RNA processing or post-transcriptional regulation (97–100). TRA2B and SRSF1 regulate pre-mRNA splicing (97, 98), whereas SYNCRIP and HNRNPD are heterogeneous nuclear ribonucleoproteins involved in mRNA processing, stability, and translational control (99, 100).

The connection of these modules after addition of first-order neighboring proteins brings KPNA2 into this context. KPNA2 encodes an importin alpha-dependent, cell-cycle-associated nuclear import receptor and may therefore reflect increased nuclear transport demand during activation of transcriptional and post-transcriptional programs (101, 102). In *Retinitis pigmentosa*, a retinal degenerative disease, KPNA2 expression has been detected in Müller cells and linked to progenitor-like, S-phase-associated states, supporting its relevance in the retinal injury and reprogramming context (103, 104). KPNA2 is therefore an attractive candidate for follow up experiments, because reduced importin alpha-dependent nuclear transport could affect the availability of transcriptional regulators required for the proliferation associated Müller cell state (78, 105). KPNA2 inhibition or knockdown could be tested for its ability to attenuate Myc, E2F, and G2M associated protein signatures (78, 106–108), reduce the abundance of cell-cycle markers such as MCM2, MCM6, and MAD2L1 (109, 110), and alter the RNA processing module represented by TRA2B, SYNCRIP, SRSF1, and HNRNPD (101). These experiments would help determine whether KPNA2 contributes functionally to the cell-cycle-associated and RNA processing signature identified here or primarily reflects increased nuclear transport demand in activated Müller cells.

Although NOLC1 occurred in several positively enriched Hallmark gene sets, it did not form a connected STRING module with the shared Myc, E2F, and G2M proteins. Its established role in nucleolar organization, rRNA transcription, and nuclear ribonucleoprotein body formation supports the interpretation that increased nucleolar and RNA-processing activity contributes to the ERU-associated Müller cell signature (111, 112).

The present reanalysis also indicated activation of interferon-associated response pathways in Müller cells derived from ERU-affected horses (113). This is in line with identification of “Interferon signaling”, “Interferon alpha/beta signaling”, “Interferon gamma signaling” via Reactome analysis that was performed during the original analysis of this dataset (6). Inflammatory stress can potentially reactivate growth control pathways that are normally silenced in adult glial cells (114). Because the dataset was generated from isolated Müller cell cultures, this signature cannot be attributed to ongoing cytokine production by infiltrating immune cells. It may instead reflect an inflammatory memory of ERU-derived Müller cells, autonomous interferon production under culture conditions, or increased sensitivity to interferon signaling after exposure to the inflammatory retinal environment (113, 115). The Hallmark Protein Secretion gene set was negatively enriched in ERU-derived Müller cells, indicating that these cells retain an inflammatory response signature despite altered secretory pathway activity (68). Future studies should therefore determine whether ERU-derived Müller cells produce interferons autonomously or respond more strongly to exogenous interferon stimulation, particularly IFNγ.

This interferon-associated signature is relevant in the context of ERU because the disease is characterized by adaptive immune activation within the eye. Retinal-antigen-responsive vitreal lymphocytes have been detected in ERU, and activated CD4^+^ T-cell phenotypes have been described in affected horses (12, 13). Previous observations also showed increased MHC class II expression on resident ocular cells, including proliferating Müller cells, in ERU (116). Thus, the proliferation-associated and interferon-responsive signature identified here may have relevance beyond glial remodeling alone. If Müller cells in the inflamed retina acquire or maintain MHC class II expression, expansion or persistence of this reactive Müller cell population could increase the number of retinal cells with potential MHC class II-mediated interactions with CD4^+^ T cells (31, 117). The present data do not demonstrate antigen presentation or T-cell activation by Müller cells, but they provide a molecular context in which interferon responsiveness, cell-cycle-associated activation, and the previously described MHC class II phenotype of ERU Müller cells can be linked.

Several aspects of this use case should be considered when interpreting the results. First, the analysis was based on a small, published dataset with three biological samples per group and was performed on processed protein-level abundance values rather than raw mass spectrometry data or peptide-level models (6). The results therefore reflect a downstream reanalysis of the available protein matrix and are not intended to replace the original raw-data-based workflow. Second, missing values were handled by left-censored imputation. Although this approach is appropriate for proteins with signals below the detection limit, imputation can influence fold changes, statistical testing, protein ranking, and downstream enrichment results (118, 119). Third, Hallmark GSEA relied on gene-symbol-based mapping of equine proteins to MSigDB gene sets (68). This enables pathway-level interpretation but does not account for equine-specific pathway curation. These considerations support the use of the present analysis as a hypothesis-generating reanalysis rather than as functional validation of the proposed mechanisms.

This reanalysis illustrates the hypothesis-generating potential of MiraProt by enabling differential abundance analysis, functional enrichment, and STRING-based association network analysis within a single workflow. In the ERU dataset, the combined interpretation of these outputs resolved a proliferation-associated and interferon-responsive Müller cell signature and highlighted candidate mechanisms involving KPNA2-associated nuclear transport, c-Myc-linked regulation, cell-cycle activation, and RNA processing. These findings are not intended to establish causal mechanisms, but they define a focused set of experimentally testable hypotheses. Thus, MiraProt extends proteomic data exploration beyond quality assessment and visualization by supporting comprehensive downstream analysis and biological prioritization from existing datasets.

## Acknowledgments

The authors gratefully acknowledge Nicole and Heinrich Veit for providing eye samples and Giulia Di Garbo and Kristin Brandt for providing ERU eyes. We acknowledge the mass spectrometry measurements performed by the Metabolomics and Proteomics Core Facility at Helmholtz Munich.

## Data Availability Statement

The underlying mass spectrometry data have been previously deposited in the PRIDE repository under accession PXD058170 and are available at https://www.ebi.ac.uk/pride/ (accessed on 8/27/2026) (6). The processed protein-level data, associated metadata, and MiraProt analysis outputs underlying the present study are provided in S1 Table. MiraProt source code is openly available under the MIT License at https://github.com/AdSchmalen/MiraProt, and version 1.0.0 is archived on Zenodo (DOI: 10.5281/zenodo.22110860) (5).

## Supporting Information

**S1 Fig.**
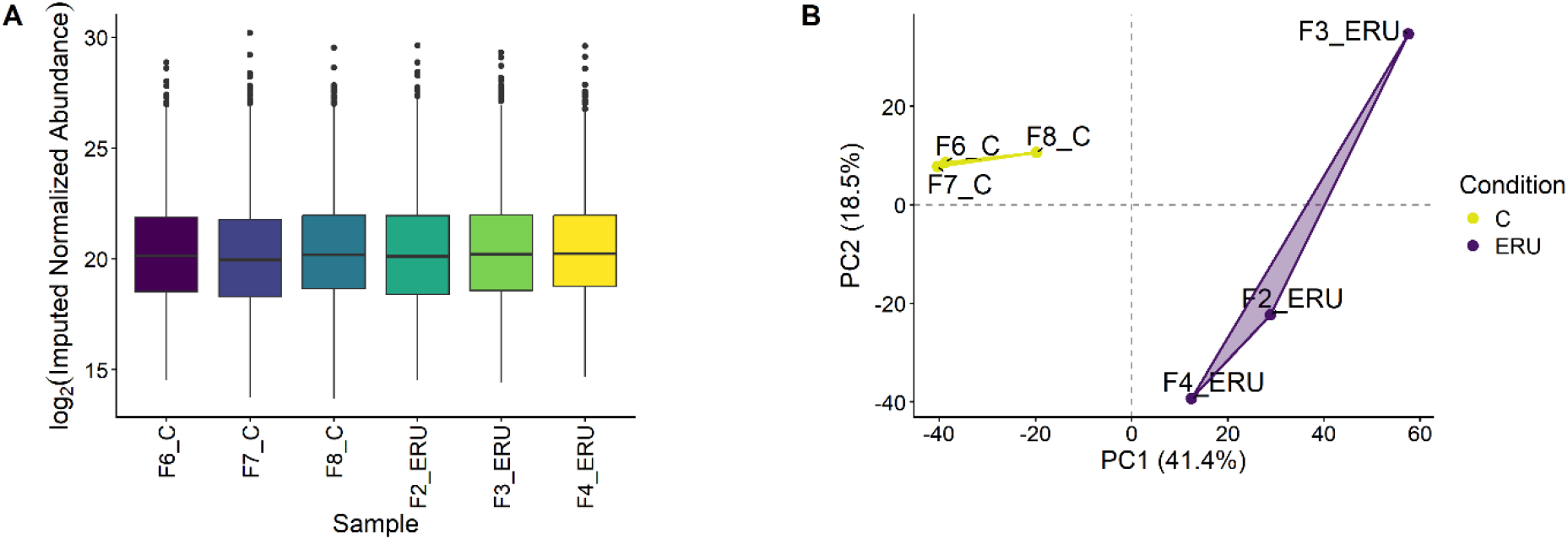
Quality control of imputed data. **(A)** Boxplot summary of imputed normalized abundance values identified in the different samples. **(B)** Principal component analysis (PCA) of the imputed data showing the two main principal components PC1 (x-axis) and PC2 (y-axis). ERU data points are depicted in purple and connected by a purple polygon and control data points C are depicted in yellow connected by a yellow polygon.

S1 Table. Comprehensive MiraProt Excel export of the ERU Müller cell proteomics analysis. The workbook contains the original and processed protein-level data, associated metadata, and the data and results generated across the MiraProt workflow, including abundance and sample-level summaries, dimensionality reduction, differential abundance visualization, functional and gene set enrichment, protein interaction analysis, protein set comparisons, and heatmap analyses.

## References

1. Chen C, Hou J, Tanner JJ, Cheng J. Bioinformatics Methods for Mass Spectrometry-Based Proteomics Data Analysis. Int J Mol Sci. 2020;21(8).

2. Jones J, MacKrell EJ, Wang TY, Lomenick B, Roukes ML, Chou TF. Tidyproteomics: an open-source R package and data object for quantitative proteomics post analysis and visualization. BMC Bioinformatics. 2023;24(1):239.

3. Lavallee-Adam M, Rauniyar N, McClatchy DB, Yates JR, 3rd. PSEA-Quant: a protein set enrichment analysis on label-free and label-based protein quantification data. J Proteome Res. 2014;13(12):5496–509.

4. Strauss MT, Bludau I, Zeng WF, Voytik E, Ammar C, Schessner JP, et al. AlphaPept: a modern and open framework for MS-based proteomics. Nat Commun. 2024;15(1):2168.

5. Schmalen A. MiraProt. 1.0.0 ed: Zenodo; 2026.

6. Fleischer AB, Amann B, von Toerne C, Degroote RL, Schmalen A, Weisser T, et al. Differential Expression of ARG1 and MRC2 in Retinal Muller Glial Cells During Autoimmune Uveitis. Biomolecules. 2025;15(2).

7. Degroote RL, Deeg CA. Immunological Insights in Equine Recurrent Uveitis. Front Immunol. 2020;11:609855.

8. Malalana F, Stylianides A, McGowan C. Equine recurrent uveitis: Human and equine perspectives. Vet J. 2015;206(1):22–9.

9. Soth R, Hoffmann ALC, Deeg CA. Enhanced ROS Production and Mitochondrial Metabolic Shifts in CD4(+) T Cells of an Autoimmune Uveitis Model. Int J Mol Sci. 2024;25(21).

10. Barfusser C, Wiedemann C, Hoffmann ALC, Hirmer S, Deeg CA. Altered Metabolic Phenotype of Immune Cells in a Spontaneous Autoimmune Uveitis Model. Front Immunol. 2021;12:601619.

11. Deeg CA, Pompetzki D, Raith AJ, Hauck SM, Amann B, Suppmann S, et al. Identification and functional validation of novel autoantigens in equine uveitis. Mol Cell Proteomics. 2006;5(8):1462–70.

12. Saldinger LK, Nelson SG, Bellone RR, Lassaline M, Mack M, Walker NJ, et al. Horses with equine recurrent uveitis have an activated CD4+ T-cell phenotype that can be modulated by mesenchymal stem cells in vitro. Vet Ophthalmol. 2020;23(1):160–70.

13. Deeg CA, Kaspers B, Gerhards H, Thurau SR, Wollanke B, Wildner G. Immune responses to retinal autoantigens and peptides in equine recurrent uveitis. Invest Ophthalmol Vis Sci. 2001;42(2):393–8.

14. Deeg CA, Ehrenhofer M, Thurau SR, Reese S, Wildner G, Kaspers B. Immunopathology of recurrent uveitis in spontaneously diseased horses. Exp Eye Res. 2002;75(2):127–33.

15. Fingerhut L, Yucel L, Strutzberg-Minder K, von Kockritz-Blickwede M, Ohnesorge B, de Buhr N. Ex Vivo and In Vitro Analysis Identify a Detrimental Impact of Neutrophil Extracellular Traps on Eye Structures in Equine Recurrent Uveitis. Front Immunol. 2022;13:830871.

16. Deeg CA, Hauck SM, Amann B, Pompetzki D, Altmann F, Raith A, et al. Equine recurrent uveitis--a spontaneous horse model of uveitis. Ophthalmic Res. 2008;40(3-4):151–3.

17. Deeg CA, Raith AJ, Amann B, Crabb JW, Thurau SR, Hauck SM, et al. CRALBP is a highly prevalent autoantigen for human autoimmune uveitis. Clin Dev Immunol. 2007;2007:39245.

18. Wang Y, Yang X, Li Q, Zhang Y, Chen L, Hong L, et al. Single-cell RNA sequencing reveals the Muller subtypes and inner blood-retinal barrier regulatory network in early diabetic retinopathy. Front Mol Neurosci. 2022;15:1048634.

19. Han C, Li Y, Zheng X, Zhang X, Li G, Zhao L, et al. AQP4- and Kir4.1-Mediated Muller Cell Oedema Is Involved in Retinal Injury Induced By Hypobaric Hypoxia. Mol Neurobiol. 2025;62(2):2012–22.

20. Eberhardt C, Amann B, Feuchtinger A, Hauck SM, Deeg CA. Differential expression of inwardly rectifying K+ channels and aquaporins 4 and 5 in autoimmune uveitis indicates misbalance in Muller glial cell-dependent ion and water homeostasis. Glia. 2011;59(5):697–707.

21. Reichenbach A, Bringmann A. New functions of Muller cells. Glia. 2013;61(5):651–78.

22. Reichenbach A, Bringmann A. Glia of the human retina. Glia. 2020;68(4):768–96.

23. Subair TI, Oyetayo B, Morales-Ramírez N, Hernández-Kelly LC, Ramírez-Martínez L, Calderón ES, et al. Glutamate Uptake Activity in Retina Müller cells: Circadian Modulation. Experimental Eye Research. 2025:110603.

24. Dun Y, Mysona B, Itagaki S, Martin-Studdard A, Ganapathy V, Smith SB. Functional and molecular analysis of D-serine transport in retinal Muller cells. Exp Eye Res. 2007;84(1):191–9.

25. Bringmann A, Pannicke T, Biedermann B, Francke M, Iandiev I, Grosche J, et al. Role of retinal glial cells in neurotransmitter uptake and metabolism. Neurochem Int. 2009;54(3-4):143–60.

26. Dyer MA, Cepko CL. Control of Muller glial cell proliferation and activation following retinal injury. Nat Neurosci. 2000;3(9):873–80.

27. Jahanshir E, Llamas J, Kim Y, Biju K, Oak S, Gnedeva K. The Hippo pathway and p27(Kip1) cooperate to suppress mitotic regeneration in the organ of Corti and the retina. Proc Natl Acad Sci U S A. 2025;122(14):e2411313122.

28. Lee SY, Surbeck JW, Drake M, Saunders A, Jin HD, Shah VA, et al. Increased Glial Fibrillary Acid Protein and Vimentin in Vitreous Fluid as a Biomarker for Proliferative Vitreoretinopathy. Invest Ophthalmol Vis Sci. 2020;61(5):22.

29. Hippert C, Graca AB, Basche M, Kalargyrou AA, Georgiadis A, Ribeiro J, et al. RNAi-mediated suppression of vimentin or glial fibrillary acidic protein prevents the establishment of Muller glial cell hypertrophy in progressive retinal degeneration. Glia. 2021;69(9):2272–90.

30. Jahnke L, Zandi S, Elhelbawi A, Conedera FM, Enzmann V. Characterization of Macroglia Response during Tissue Repair in a Laser-Induced Model of Retinal Degeneration. Int J Mol Sci. 2023;24(11).

31. Schmalen A, Lorenz L, Grosche A, Pauly D, Deeg CA, Hauck SM. Proteomic Phenotyping of Stimulated Muller Cells Uncovers Profound Pro-Inflammatory Signaling and Antigen-Presenting Capacity. Front Pharmacol. 2021;12:771571.

32. Eastlake K, Banerjee PJ, Angbohang A, Charteris DG, Khaw PT, Limb GA. Muller glia as an important source of cytokines and inflammatory factors present in the gliotic retina during proliferative vitreoretinopathy. Glia. 2016;64(4):495–506.

33. Vecino E, Rodriguez FD, Ruzafa N, Pereiro X, Sharma SC. Glia-neuron interactions in the mammalian retina. Prog Retin Eye Res. 2016;51:1–40.

34. Zhu Y, Orre LM, Zhou Tran Y, Mermelekas G, Johansson HJ, Malyutina A, et al. DEqMS: A Method for Accurate Variance Estimation in Differential Protein Expression Analysis. Mol Cell Proteomics. 2020;19(6):1047–57.

35. Chang W, Cheng J, Allaire J, Sievert C, Schloerke B, Xie Y, et al. shiny: Web Application Framework for R. 1.10.0 ed2024.

36. Attali D. shinyjs: Easily Improve the User Experience of Your Shiny Apps in Seconds. 2.1.0 ed2021.

37. Perrier V, Meyer F, Granjon D. shinyWidgets: Custom Inputs Widgets for Shiny. 0.8.7 ed2024.

38. Wickham H. ggplot2: Elegant Graphics for Data Analysis. 3.5.1 ed: Springer-Verlag New York; 2016.

39. Sievert C. Interactive Web-Based Data Visualization with R, plotly, and shiny. 4.10.4 ed: Chapman and Hall/CRC; 2020.

40. Huynh T, Ramachandran G, Banerjee S, Monteiro J, Stenzel M, Sandler DP, et al. Comparison of methods for analyzing left-censored occupational exposure data. Ann Occup Hyg. 2014;58(9):1126–42.

41. Stekhoven DJ, Buhlmann P. MissForest--non-parametric missing value imputation for mixed-type data. Bioinformatics. 2012;28(1):112–8.

42. van Buuren S, Groothuis-Oudshoorn K. mice: Multivariate Imputation by Chained Equations in R. J Stat Softw. 2011;45(3):1–67.

43. Brown MB, Forsythe AB. The Small Sample Behavior of Some Statistics Which Test the Equality of Several Means. Technometrics. 1974;16:129–32.

44. Zimmerman DW, Zumbo BD. Rank transformations and the power of the Student t test and Welch t’ test for non-normal populations with unequal variances. Canadian Journal of Experimental Psychology / Revue canadienne de psychologie expérimentale. 1993;47(3):523–39.

45. Smyth GK. Linear models and empirical bayes methods for assessing differential expression in microarray experiments. Stat Appl Genet Mol Biol. 2004;3:Article3.

46. Mann HB, Whitney DR. On a Test of Whether one of Two Random Variables is Stochastically Larger than the Other. The Annals of Mathematical Statistics. 1947;18(1):50–60, 11.

47. Dunn OJ. Multiple Comparisons Among Means. Journal of the American Statistical Association. 1961;56(293):52–64.

48. Holm S. A Simple Sequentially Rejective Multiple Test Procedure. Scandinavian Journal of Statistics. 1979;6(2):65–70.

49. Hochberg Y. A Sharper Bonferroni Procedure for Multiple Tests of Significance. Biometrika. 1988;75(4):800–2.

50. Hommel G. A Stagewise Rejective Multiple Test Procedure Based on a Modified Bonferroni Test. Biometrika. 1988;75(2):383–6.

51. Benjamini Y, Hochberg Y. Controlling the False Discovery Rate: A Practical and Powerful Approach to Multiple Testing. Journal of the Royal Statistical Society Series B (Methodological). 1995;57(1):289–300.

52. Benjamini Y, Yekutieli D. The control of the false discovery rate in multiple testing under dependency. The Annals of Statistics. 2001;29(4):1165–88, 24.

53. Huber W, Carey VJ, Gentleman R, Anders S, Carlson M, Carvalho BS, et al. Orchestrating high-throughput genomic analysis with Bioconductor. Nat Methods. 2015;12(2):115–21.

54. Durinck S, Spellman PT, Birney E, Huber W. Mapping identifiers for the integration of genomic datasets with the R/Bioconductor package biomaRt. Nat Protoc. 2009;4(8):1184–91.

55. Ma S, Dai Y. Principal component analysis based methods in bioinformatics studies. Brief Bioinform. 2011;12(6):714–22.

56. Shah N, Meng Q, Zou Z, Zhang X. Systematic analysis on the horse-shoe-like effect in PCA plots of scRNA-seq data. Bioinform Adv. 2024;4(1):vbae109.

57. Becht E, McInnes L, Healy J, Dutertre CA, Kwok IWH, Ng LG, et al. Dimensionality reduction for visualizing single-cell data using UMAP. Nat Biotechnol. 2018.

58. Robinson MD, McCarthy DJ, Smyth GK. edgeR: a Bioconductor package for differential expression analysis of digital gene expression data. Bioinformatics. 2010;26(1):139–40.

59. Yu G. Thirteen years of clusterProfiler. Innovation (Camb). 2024;5(6):100722.

60. Xu S, Hu E, Cai Y, Xie Z, Luo X, Zhan L, et al. Using clusterProfiler to characterize multiomics data. Nat Protoc. 2024;19(11):3292–320.

61. Wu T, Hu E, Xu S, Chen M, Guo P, Dai Z, et al. clusterProfiler 4.0: A universal enrichment tool for interpreting omics data. Innovation (Camb). 2021;2(3):100141.

62. Yu G, Wang LG, Han Y, He QY. clusterProfiler: an R package for comparing biological themes among gene clusters. OMICS. 2012;16(5):284–7.

63. Subramanian A, Tamayo P, Mootha VK, Mukherjee S, Ebert BL, Gillette MA, et al. Gene set enrichment analysis: a knowledge-based approach for interpreting genome-wide expression profiles. Proc Natl Acad Sci U S A. 2005;102(43):15545–50.

64. Mootha VK, Lindgren CM, Eriksson KF, Subramanian A, Sihag S, Lehar J, et al. PGC-1alpha-responsive genes involved in oxidative phosphorylation are coordinately downregulated in human diabetes. Nat Genet. 2003;34(3):267–73.

65. Zyla J, Marczyk M, Weiner J, Polanska J. Ranking metrics in gene set enrichment analysis: do they matter? BMC Bioinformatics. 2017;18(1):256.

66. Korotkevich G, Sukhov V, Budin N, Shpak B, Artyomov MN, Sergushichev A. Fast gene set enrichment analysis. bioRxiv. 2021:060012.

67. Tarca AL, Draghici S, Bhatti G, Romero R. Down-weighting overlapping genes improves gene set analysis. BMC Bioinformatics. 2012;13:136.

68. Liberzon A, Birger C, Thorvaldsdottir H, Ghandi M, Mesirov JP, Tamayo P. The Molecular Signatures Database (MSigDB) hallmark gene set collection. Cell Syst. 2015;1(6):417–25.

69. Szklarczyk D, Kirsch R, Koutrouli M, Nastou K, Mehryary F, Hachilif R, et al. The STRING database in 2023: protein-protein association networks and functional enrichment analyses for any sequenced genome of interest. Nucleic Acids Res. 2023;51(D1):D638–D46.

70. B.V. A, Thieurmel B. visNetwork: Network Visualization using ’vis.js’ Library. 2.1.2 ed2022.

71. Girvan M, Newman ME. Community structure in social and biological networks. Proc Natl Acad Sci U S A. 2002;99(12):7821–6.

72. Csárdi G, Nepusz T, editors. The igraph software package for complex network research 2006.

73. Csárdi G, Nepusz T, Traag V, Horvát S, Zanini F, Noom D, et al. igraph: Network Analysis and Visualization in R. 2025.

74. Lex A, Gehlenborg N, Strobelt H, Vuillemot R, Pfister H. UpSet: Visualization of Intersecting Sets. IEEE Trans Vis Comput Graph. 2014;20(12):1983–92.

75. Chen H. VennDiagram: Generate High-Resolution Venn and Euler Plots. 1.7.3 ed 2022.

76. Krassowski M. ComplexUpset. 1.3.3 ed 2020.

77. Gu Z, Eils R, Schlesner M. Complex heatmaps reveal patterns and correlations in multidimensional genomic data. Bioinformatics. 2016;32(18):2847–9.

78. Lee MS, Jui J, Sahu A, Goldman D. Mycb and Mych stimulate Muller glial cell reprogramming and proliferation in the uninjured and injured zebrafish retina. Development. 2024;151(14).

79. Zhu W, Giangrande PH, Nevins JR. E2Fs link the control of G1/S and G2/M transcription. EMBO J. 2004;23(23):4615–26.

80. Ritchie ME, Phipson B, Wu D, Hu Y, Law CW, Shi W, et al. limma powers differential expression analyses for RNA-sequencing and microarray studies. Nucleic Acids Res. 2015;43(7):e47.

81. Leek JT, Johnson WE, Parker HS, Jaffe AE, Storey JD. The sva package for removing batch effects and other unwanted variation in high-throughput experiments. Bioinformatics. 2012;28(6):882–3.

82. Chen X, Zhang B, Wang T, Bonni A, Zhao G. Robust principal component analysis for accurate outlier sample detection in RNA-Seq data. BMC Bioinformatics. 2020;21(1):269.

83. Dorrity MW, Saunders LM, Queitsch C, Fields S, Trapnell C. Dimensionality reduction by UMAP to visualize physical and genetic interactions. Nat Commun. 2020;11(1):1537.

84. Peng H, Wang H, Kong W, Li J, Goh WWB. Optimizing differential expression analysis for proteomics data via high-performing rules and ensemble inference. Nat Commun. 2024;15(1):3922.

85. Sticker A, Goeminne L, Martens L, Clement L. Robust Summarization and Inference in Proteome-wide Label-free Quantification. Mol Cell Proteomics. 2020;19(7):1209–19.

86. Killinger BJ, Petyuk VA, Wright AT. Detecting differential protein abundance by combining peptide level P-values. Mol Omics. 2020;16(6):554–62.

87. Choi H, Kim S, Fermin D, Tsou CC, Nesvizhskii AI. QPROT: Statistical method for testing differential expression using protein-level intensity data in label-free quantitative proteomics. J Proteomics. 2015;129:121–6.

88. Wieczorek S, Combes F, Borges H, Burger T. Protein-Level Statistical Analysis of Quantitative Label-Free Proteomics Data with ProStaR. Methods Mol Biol. 2019;1959:225–46.

89. Liberzon A, Subramanian A, Pinchback R, Thorvaldsdottir H, Tamayo P, Mesirov JP. Molecular signatures database (MSigDB) 3.0. Bioinformatics. 2011;27(12):1739–40.

90. Coller HA, Grandori C, Tamayo P, Colbert T, Lander ES, Eisenman RN, et al. Expression analysis with oligonucleotide microarrays reveals that MYC regulates genes involved in growth, cell cycle, signaling, and adhesion. Proc Natl Acad Sci U S A. 2000;97(7):3260–5.

91. Popay TM, Wang J, Adams CM, Howard GC, Codreanu SG, Sherrod SD, et al. MYC regulates ribosome biogenesis and mitochondrial gene expression programs through its interaction with host cell factor-1. Elife. 2021;10.

92. Ren B, Cam H, Takahashi Y, Volkert T, Terragni J, Young RA, et al. E2F integrates cell cycle progression with DNA repair, replication, and G(2)/M checkpoints. Genes Dev. 2002;16(2):245–56.

93. Ishida S, Huang E, Zuzan H, Spang R, Leone G, West M, et al. Role for E2F in control of both DNA replication and mitotic functions as revealed from DNA microarray analysis. Mol Cell Biol. 2001;21(14):4684–99.

94. Saldivar JC, Hamperl S, Bocek MJ, Chung M, Bass TE, Cisneros-Soberanis F, et al. An intrinsic S/G(2) checkpoint enforced by ATR. Science. 2018;361(6404):806–10.

95. Saito Y, Santosa V, Ishiguro KI, Kanemaki MT. MCMBP promotes the assembly of the MCM2-7 hetero-hexamer to ensure robust DNA replication in human cells. Elife. 2022;11.

96. Lu X, Zhang Y, Xue J, Evert M, Calvisi D, Chen X, et al. MAD2L1 supports MYC-driven liver carcinogenesis in mice and predicts poor prognosis in human hepatocarcinoma. Toxicol Sci. 2025;203(1):41–51.

97. Zhou T, Fu P, Chen D, Liu R. Transformer 2beta regulates the alternative splicing of cell cycle regulatory genes to promote the malignant phenotype of ovarian cancer. Oncol Res. 2023;31(5):769–85.

98. Wu Q, Yu H, Sun H, Lv J, Zhuang J, Cai L, et al. SRSF1-mediated alternative splicing regulates bladder cancer progression and cisplatin sensitivity through HIF1A/BNIP3/mitophagy axis. J Transl Med. 2025;23(1):571.

99. Li C, Lu T, Chen H, Yu Z, Chen C. The up-regulation of SYNCRIP promotes the proliferation and tumorigenesis via DNMT3A/p16 in colorectal cancer. Sci Rep. 2024;14(1):21570.

100. Hu H, Zhang H, Xing Y, Zhou Y, Chen J, Li C, et al. The lncRNA THOR interacts with and stabilizes hnRNPD to promote cell proliferation and metastasis in breast cancer. Oncogene. 2022;41(49):5298–314.

101. Cao L, Jia K, Van Tine BA, Yu Y, Peng Y, Chen X, et al. KPNA2 promotes osteosarcoma progression by regulating the alternative splicing of DDX3X mediated by YBX1. Oncogene. 2025;44(26):2186–200.

102. Chen X, Wei H, Yue A, Zhang H, Zheng Y, Sun W, et al. KPNA2 promotes the progression of gastric cancer by regulating the alternative splicing of related genes. Sci Rep. 2024;14(1):17140.

103. Roesch K, Stadler MB, Cepko CL. Gene expression changes within Muller glial cells in retinitis pigmentosa. Mol Vis. 2012;18:1197–214.

104. Trimarchi JM, Stadler MB, Cepko CL. Individual retinal progenitor cells display extensive heterogeneity of gene expression. PLoS One. 2008;3(2):e1588.

105. Wang Y, Jin Y, Li X. A positive feedback loop between KPNA2 and FOXM1 promotes the proliferation of lung adenocarcinoma. Eur J Med Res. 2025;31(1):122.

106. Mitra S, Sharma P, Kaur S, Khursheed MA, Gupta S, Chaudhary M, et al. Dual regulation of lin28a by Myc is necessary during zebrafish retina regeneration. J Cell Biol. 2019;218(2):489–507.

107. Xiang S, Wang Z, Ye Y, Zhang F, Li H, Yang Y, et al. E2F1 and E2F7 differentially regulate KPNA2 to promote the development of gallbladder cancer. Oncogene. 2019;38(8):1269–81.

108. Huang L, Wang HY, Li JD, Wang JH, Zhou Y, Luo RZ, et al. KPNA2 promotes cell proliferation and tumorigenicity in epithelial ovarian carcinoma through upregulation of c-Myc and downregulation of FOXO3a. Cell Death Dis. 2013;4(8):e745.

109. Ohtani K, Iwanaga R, Nakamura M, Ikeda M, Yabuta N, Tsuruga H, et al. Cell growth-regulated expression of mammalian MCM5 and MCM6 genes mediated by the transcription factor E2F. Oncogene. 1999;18(14):2299–309.

110. Hernando E, Nahle Z, Juan G, Diaz-Rodriguez E, Alaminos M, Hemann M, et al. Rb inactivation promotes genomic instability by uncoupling cell cycle progression from mitotic control. Nature. 2004;430(7001):797–802.

111. Chen HK, Pai CY, Huang JY, Yeh NH. Human Nopp140, which interacts with RNA polymerase I: implications for rRNA gene transcription and nucleolar structural organization. Mol Cell Biol. 1999;19(12):8536–46.

112. Bizarro J, Deryusheva S, Wacheul L, Gupta V, Ernst FGM, Lafontaine DLJ, et al. Nopp140-chaperoned 2’-O-methylation of small nuclear RNAs in Cajal bodies ensures splicing fidelity. Genes Dev. 2021;35(15-16):1123–41.

113. Hauck SM, Schoeffmann S, Amann B, Stangassinger M, Gerhards H, Ueffing M, et al. Retinal Mueller glial cells trigger the hallmark inflammatory process in autoimmune uveitis. J Proteome Res. 2007;6(6):2121–31.

114. Iribarne M, Hyde DR, Masai I. TNFalpha Induces Muller Glia to Transition From Non-proliferative Gliosis to a Regenerative Response in Mutant Zebrafish Presenting Chronic Photoreceptor Degeneration. Front Cell Dev Biol. 2019;7:296.

115. Drescher KM, Whittum-Hudson JA. Evidence for induction of interferon-alpha and interferon-beta in retinal glial cells of Muller. Virology. 1997;234(2):309–16.

116. Romeike A, Brugmann M, Drommer W. Immunohistochemical studies in equine recurrent uveitis (ERU). Vet Pathol. 1998;35(6):515–26.

117. Quinn J, Salman A, Paluch C, Jackson-Wood M, McClements ME, Luo J, et al. Single-cell transcriptomic analysis of retinal immune regulation and blood-retinal barrier function during experimental autoimmune uveitis. Sci Rep. 2024;14(1):20033.

118. Liu M, Dongre A. Proper imputation of missing values in proteomics datasets for differential expression analysis. Brief Bioinform. 2021;22(3).

119. Harris L, Fondrie WE, Oh S, Noble WS. Evaluating Proteomics Imputation Methods with Improved Criteria. J Proteome Res. 2023;22(11):3427–38.

